# A dual foveal-peripheral visual processing model implements efficient saccade selection

**DOI:** 10.1101/725879

**Authors:** Emmanuel Daucé, Pierre Albiges, Laurent U Perrinet

**Affiliations:** Institut de Neurosciences de la Timone, CNRS/Aix-Marseille Université, France

## Abstract

We develop a visuomotor model that implements visual search as a focal accuracy-seeking policy, with the target’s position and category drawn independently from a common generative process. Consistently with the anatomical separation between the ventral versus dorsal pathways, the model is composed of two pathways, that respectively infer what to see and where to look. The “What” network is a classical deep learning classifier, that only processes a small region around the center of fixation, providing a “foveal” accuracy. In contrast, the “Where” network processes the full visual field in a biomimetic fashion, using a log-polar retinotopic encoding, which is preserved up to the action selection level. The foveal accuracy is used to train the “Where” network. After training, the “Where” network provides an “accuracy map” that serves to guide the eye toward peripheral objects. The comparison of both networks accuracies amounts to either select a saccade or to keep the eye at the center to identify the target. We test this setup on a simple task of finding a digit in a large, cluttered image. Our simulation results demonstrate the effectiveness of this approach, increasing by one order of magnitude the radius of the visual field toward which the agent can detect and recognize a target, either through a single saccade or with multiple ones. Importantly, our log-polar treatment of the visual information exploits the strong compression rate performed at the sensory level, providing ways to implement visual search in a sub-linear fashion, in contrast with mainstream computer vision.

## 1 Introduction

### 1.1 Problem statement

The field of computer vision was recently recast by the outstanding capability of convolution-based deep neural networks to capture the semantic content of images and photographs. Human performance is now outreached by computer algorithms in numerous image categorization tasks (He et al., 2015). One of the reasons explaining this breakthrough is a strong reduction in the number of parameters used to train the network, through a massive sharing of weights in the convolutional layers. Reducing the number of parameters and/or the size of the visual data that needs to be processed is a decisive factor for further improvements. Initially trained on energy greedy, high-performance computers, these algorithms are now designed to work on more common hardware such as desktop computers with dedicated GPU hardware (Sandler et al., 2018). Despite lots of efforts both in hardware and software optimization, the processing of pixel-based images is still done at a cost that scales linearly with the image size: All pixels present in the image are systematically processed by the computer algorithm, even the ones that are useless for the task at hand. Current computer vision algorithms consequently manipulate millions of pixels and millions of variables with ensuing energy consumption, even in the case of downsampled images, and with a still prohibitive cost for large images and videos. The need to detect visual objects at a glance while running on resource-constrained embedded hardware, for instance in autonomous driving, introduces a necessary trade-off between efficiency and accuracy, requiring renewed mathematical treatment and computational implementations.

Interestingly, things work differently when human vision is considered. First, human vision is still unsurpassable in the case of ecological real-time sensory flows. Indeed, object recognition can be achieved by the human visual system both rapidly, – in less than 100 ms (Kirchner and Thorpe, 2006) – and at a low energy cost (< 5 *W*). On top of that, it is mostly self-organized, robust to visual transforms or lighting conditions and can learn with few examples. If many different anatomical features may explain this efficiency, the main difference lies in the fact that its sensor (the retina) combines a non-homogeneous sampling of the world with the capacity to rapidly change its center of fixation: on the one hand, the retina is composed of two separate systems: a central, high definition fovea (a disk of about 6 degrees of diameter in visual angle around the center of gaze) and a large, lower definition peripheral area (Strasburger et al., 2011). On the other hand, the human vision is active and dynamic: the retina is attached at the back of the eye which is capable of low latency, high-speed eye movements. In particular, saccades are stereotyped eye movements that allow for efficient changes of the position of the center of gaze: they take about 200 ms to initiate, last about 200 ms and usually reach a maximum velocity of approx 600 degrees per second (Bahill et al., 1975). The scanning of a full visual scene is thus not done in parallel but sequentially, and only scene-relevant regions of interest are scanned through saccades. This implies a *decision process* between each saccade that decides *where to look next*. This behaviour is prevalent in biological vision with on average a saccade every 2 seconds, that is, almost a billion saccade in a lifetime. The interplay of peripheral search and focal inspection allows human observers to engage in an integrated action/perception loop that sequentially scans and analyses the different parts of the visual scene.

Take for instance the case of an encounter with a friend in a crowded café. To catch the moment of his/her arrival, a face-seeking visual search is needed, possibly under heavy sensory clutter conditions. To do so, relevant parts of the visual scene need to be scanned sequentially with the gaze. Each saccade may potentially allow you to recognize your friend, provided it is accurately focused on each target faces. The main feature of this task is thus the monitoring of a particular *class* of objects (e.g. human faces) in the periphery of the visual field before the actual eye displacement, and the processing of the foveal visual data. Searching for *any* face in a peripheral and crowded display needs thus to precede the recognition of a specific face *identity*. For the biological vision is the result of a continual optimization under strong material and energy constraints via natural selection, it is important to understand both its ground principles and its specific computational and material constraints in order to implement effective biomimetic vision systems. The problem we address is thus how to ground an artificial visual processing system on top of the material constraints found in human vision, that is conforming to the structure of the visual input and to the capability of the visual apparatus to rapidly scan a visual scene through saccades, in order to find and identify objects of interest. We thus start from an elementary visual search problem, which is how to locate an object in a large, cluttered image, and take human vision as a guide for efficient design.

### 1.2 State of the art

The visual search problem, that is finding and identifying objects in a visual scene, is a classical task in computer vision, appealing as well to machine learning, signal processing or robotics. Crucially, it also speaks to neuroscience, for it refers to the mechanisms underlying foveation and more generally to low-level attention mechanisms. When restricted to a mere “feature search” (Treisman and Gelade, 1980), many computational solutions are proposed in the computer vision literature. Notably, recent advances in deep learning have been proven efficient to solve the task with models such as faster-RCNN (Ren et al., 2017) or YOLO (Redmon et al., 2016). Typical object search implementations predict in the image the probability of proposed bounding boxes around visual objects. While rapid, the potential number of boxes may significantly increase with image size and the approach more generally necessitates dedicated hardware to run in real time (Feng et al., 2019). Under fine-tailored algorithmic and material optimization, the visual search problem can be considered in the best case as *linear* in the number of pixels (Strengert et al., 2006), which still represents a heavy load for real-time image processing. This poses the problem of the *energy scaling* of current computer vision algorithms to large/high definition visual displays. This scaling problem becomes even more crucial when considering a dynamical stream of sensory images.

Analogously to human visual search strategies, low-level attentional mechanisms may help guide the localization of targets. A sequence of saccades over a natural scene defines a scan-path which provides ways to define *saliency maps* (Itti and Koch, 2001). These quantify the attractiveness of the different parts of an image that are consistent with the detection of objects of interest. Essential to understand and predict saccades, they also serve as phenomenological models of attention. Estimating the saliency map from a luminous image is a classical problem in neuroscience, that was shown to be consistent with a distance from baseline image statistics known as the “Bayesian surprise” (Itti and Baldi, 2009). Such an approach was extended in the AIM (Bruce and Tsotsos, 2009) and SUN (Zhang et al., 2008) models. Recently, the saliency approach was updated using deep learning to estimate saliency maps over large databases of natural images (Kummerer et al., 2017). While efficient at predicting the probability of fixation, these methods miss an essential component in the action-perception loop: they operate on the full image while the retina operates on the non-uniform, foveated sampling of visual space (see Figure 1-C). Herein, we believe that this constitutes an essential factor to reproduce and understand the active vision process.

**Figure 1.**
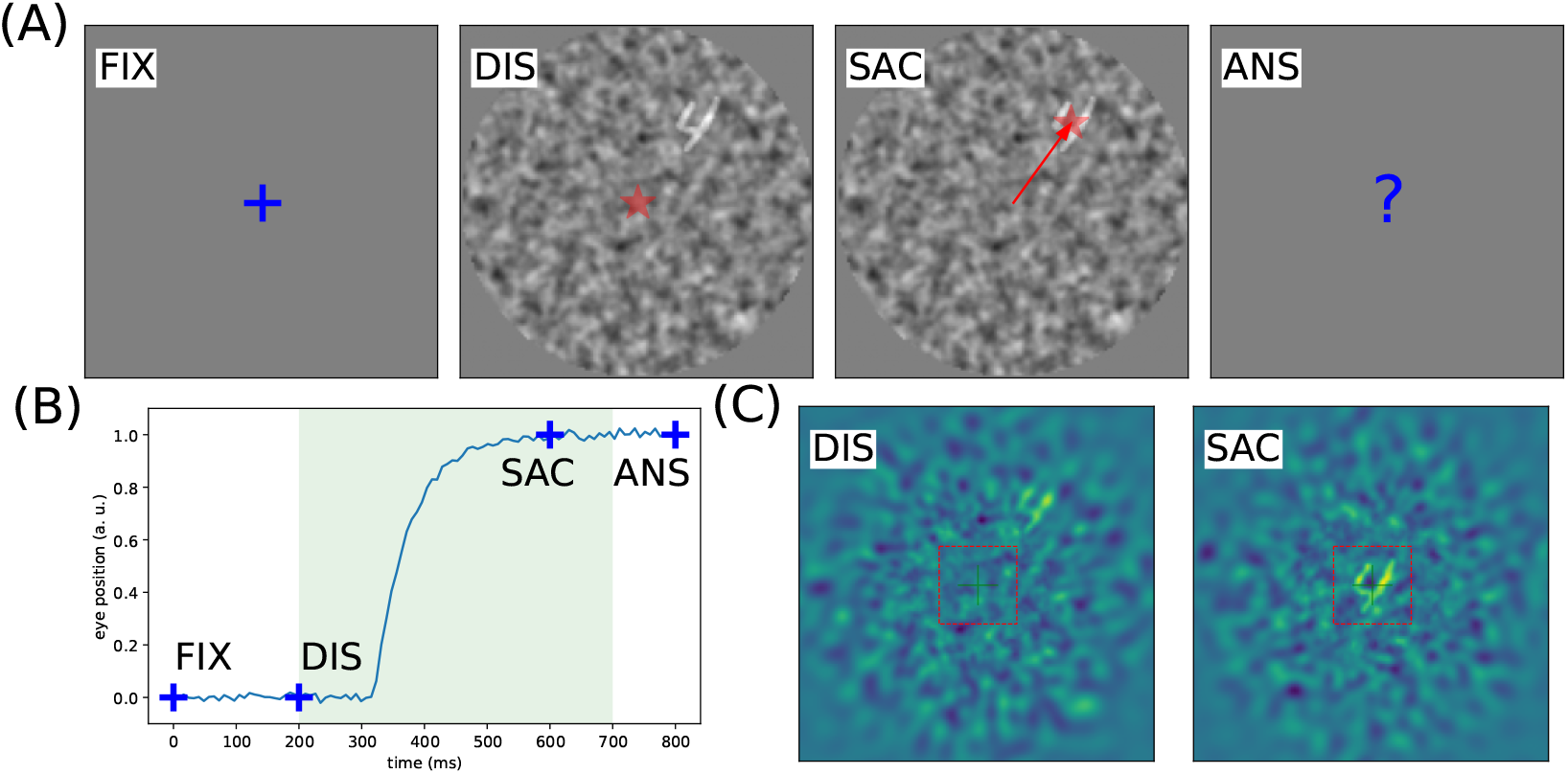
Problem setting: In generic, ecological settings, when searching for one target (from a class of targets) in a cluttered environment, the visual system is bound with an action selection problem. It is synthesized in the following virtual experiment: **(A)** After a fixation period `FIX` of 200 ms, an observer is presented with a luminous display `DIS` showing a single target from a known class (here digits) put at a random position within the field of view. The display is presented for a short period of 500 ms (light shaded area in B), which is enough to perform at most one saccade on the potential target (`SAC`, here successful). Finally, the observer has to identify the digit by a keypress `ANS`. *NB*: the target contrast is here enhanced to 100% for better readability. **(B)** Prototypical trace of a saccadic eye movement to the target position. In particular, we show the fixation window `FIX` and the temporal window during which a saccade is possible (green shaded area). **(C)** Simulated reconstruction of the visual information from the internal retinotopic map at the onset of the display `DIS` and after a saccade `SAC`, the dashed red box indicating the foveal region. The task does not consist in inferring the location of the target, but rather to infer an action that may provide relevant pixels at the center of fixation, allowing to identify the target’s category. By comparison with the external display (see A), the action is processed from log-polar coefficients, representing a focal sample of the total visual field. Controlling the clutter and reducing the contrast of the digit allows to modulate the task’s difficulty.

Foveated models of vision have been considered for a long time in robotics and computer vision as a way to leverage the visual scene scaling problem. Focal computer vision relies on a non-homogeneous compression of an image, that maintains the pixel information at the center of fixation and strongly compresses it at the periphery, including pyramidal encoding (Butko and Movellan, 2010, Kortum and Geisler, 1996), local wavelet decomposition (Daucé, 2018) and log-polar encoding (Javier Traver and Bernardino, 2010). A recent deep-learning-based implementation of such compression shows that in a video flow, a log-polar sampling of the image is sufficient to provide a reconstruction of the whole image (Kaplanyan et al., 2019). However, this particular algorithm lacks a system predicting the best saccadic action to perform. In summary, though focal and multi-scale encoding is now largely considered in static computer vision, sequential implementations have not been shown effective enough to overtake static object search methods. Several implementations of a focal sequential search in visual processing can be found in the literature, with various degrees of biological realism (Fu et al., 2017, Mnih et al., 2014), that often rely on a simplified focal encoding, long training procedures and bounded sequential processing. More realistic attempts to combine foveal encoding and sequential visual search can be found in (Butko and Movellan, 2010, Denil et al., 2012), to which our approach is compared later on.

In contrast with the phenomenological (or “bottom-up”) approaches, active models of vision (Butko and Movellan, 2010, Daucé, 2018, Najemnik and Geisler, 2005) provide the ground principles of saccadic exploration. In general, they assume the existence of a generative model from which both the target position and category can be inferred through active sampling. This comes from the constraint that the visual sensor is foveated but can generate a saccade. Several studies are relevant to our endeavor. First, one can consider optimal strategies to solve the problem of the visual search of a target (Najemnik and Geisler, 2005). In a setting similar to that presented in Figure 1-A, where the target is an oriented edge and the background is defined as pink noise, authors show first that a Bayesian ideal observer comes out with an optimal strategy, and second that human observers are close to that optimal performance. Though appropriately predicting sequences of saccades in a perception action loop, this model is limited by the simplicity of the display (elementary edges added on stationary noise, a finite number of non-overlapping locations on a discrete grid) and by the abstract level of modeling. Despite these (inevitable) simplifications, this study could successfully predict some key characteristics of visual scanning such as the trade-off between memory content and speed. Looking more closely at neurophysiology, the study of (Samonds et al., 2018) allows to go further in understanding the interplay between saccadic behavior and the statistics of the input. In this study, authors were able to manipulate the size of saccades by monitoring key properties of the presented (natural) images. For instance, smaller images generate smaller saccades. Interestingly, they also predicted the size of saccades from the size of visual receptive fields for different species, including mice which lack a foveal region. One key prediction of this study which is relevant for our problem is the fact that saccades seem optimal to *a priori* decorrelate the visual input, that is, to minimize redundancy in the sequence of generated saccades, knowing the statistics of the visual inputs.

A further modeling perspective is provided by (Friston et al., 2012). In this setup, a full description of the visual world is used as a generative process. An agent is completely described by the generative model governing the dynamics of its internal beliefs and is interacting with this image by scanning it through a foveated sensor, just as described in Figure 1. Thus, equipping the agent with the ability to actively sample the visual world allows to interpret saccades as optimal experiments, by which the agent seeks to confirm predictive models of the (hidden) world. One key ingredient to this process is the (internal) representation of counterfactual predictions, that is, the probable consequences of possible hypothesis as they would be realized into actions (here, saccades). Following such an active inference scheme, numerical simulations reproduce sequences of eye movements that fit well with empirical data (Mirza et al., 2018). Compared to previous studies as (Najemnik and Geisler, 2005), the saccades are not the output of a value-based cost function, such as a saliency map, but are the result of minimizing the entropy of the agent’s internal beliefs, knowing the generative model of the visual world.

### 1.3 Outline

So far, few models in active vision come with an integrated processing of the visual scene, from early visual treatment toward saccade selection. The difficulty lies in combining object hypothesis (feature) space along with their spatial mapping. As pointed out earlier, the system needs to guess where the interesting objects lie in space before actually knowing what they are. Establishing the position of the objects in space is thus crucial, for it resorts to the capability of the eye to reach them with a saccade, so as to finally identify them. Inferring target’s position in the peripheral visual field is thus an essential component of focal visual processing, and the acuity of such target selection ultimately conditions the capability to rapidly and efficiently process the scene. Stemming from the active vision principles, we thus address the question of the interplay of the location and identity processing in vision, and provide an artificial vision setup that efficiently implements those principles. Herein, our framework is made as general as possible, with minimal mathematical treatment, to speak largely to fragmented domains, such as machine learning, neuroscience and robotics.

The paper is organized as follows. After this introduction, the principles underlying accuracy-based saccadic control are defined in the second section. We first define notations, variables and equations for the generative process governing the experiment and the generative model for the active vision agent. Complex combinatorial inferences are here replaced by separate pathways, i.e. the spatial (“Where”) and categorical (“What”) pathways, whose output is combined to infer optimal eye displacements and subsequent identification of the target. Our agent, equipped with a foveated sensor, should learn an optimal behavior strategy to actively scan the visual scene. Numerical simulations are presented in the results section, demonstrating the applicability of this framework to tasks with different complexity levels. The discussion section finally summarizes the results, showing its relative advantages in comparison with other frameworks, and providing ways toward possible improvements. Implementation details are provided in the methods section, giving ways to reproduce our results, showing in particular how to simplify the learning using accuracy-driven action maps.

## 2 Setup

### 2.1 Experimental design

In order to implement our visual processing setup, we provide a simplified visual environment toward which a visual agent can act on. This visual search task is formalized and simplified in a way reminiscent to classical psychophysical experimentation: an observer is asked to classify digits (for instance as taken from the MNIST dataset, as introduced by (Lecun et al., 1998)) as they are shown with a given size on a computer display. However, these digits can be placed at random positions on the display, and visual clutter is added as a background to the image (see Figure 1-A). In order to vary the difficulty of the task, different parameters are controlled, such as the target eccentricity, the background noise period and and the signal/noise ratio (SNR). The agent initially fixates the center of the screen. Due to the peripheral clutter, he needs to explore the visual scene through saccades to provide the answer. He controls a foveal visual sensor that can move over the visual scene through saccades (see Figure 1-B). When a saccade is actuated, the center of fixation moves toward a new location, which updates the visual input (see Figure 1-C). The lower the SNR and the larger the initial target eccentricity, the more difficult the identification. There is a range of eccentricities for which it is impossible to identify the target from a single glance, so that a saccade is necessary to reduce the relative eccentricity and issue a proper response. This setup implies also that the position of the object may be detected in the peripheral clutter *before* being properly identified.

This setup provides the conditions for a separate processing of the visual information. On the one side, the detailed information present at the center of fixation needs to be analyzed to provide specific environmental cues. On the other side, the full visual field, i.e. mainly the low resolution part surrounding the fovea, needs to be processed in order to identify regions of interest that deserve fixation. This basically means making a choice of “what’s interesting next”. The actual content of putative peripheral locations does not need to be known in advance, but it needs to look interesting enough, and of course to be reachable by a saccade. This is reminiscent of the What/Where visual processing separation observed in primates’ ventral and dorsal visual pathways (Mishkin et al., 1983).

### 2.2 Computational implementation

In order to show it is possible to learn such a task, it is sufficient to demonstrate the existence of a simple “Deep Learning” neural network that would implement it, for instance through the effective success of its training. More specifically, this class of modern parametric classifiers are composed of many layers (hence the terminology) that can be trained through gradient descent over arbitrary input and output feature spaces. For our specific problem, the anatomy of the agent is made of two separate pathways for which a different processing is realized by two different neural networks (see Figure 2). The proposed computational architecture is connected in a closed-loop fashion to the visual environment, with the capacity to produce saccades whose effect is to shift the visual field from one visual position to another. By analogy with biological vision, the identification of the target is assumed to rely on the very central part of the retina (the fovea), that comes with higher density of cones, and thus higher spatial precision. In contrast, the saccade planning should rely on the full visual field, with peripheral regions having a lower density of sensors and thus a lower sensitivity to high spatial frequencies.

**Figure 2.**
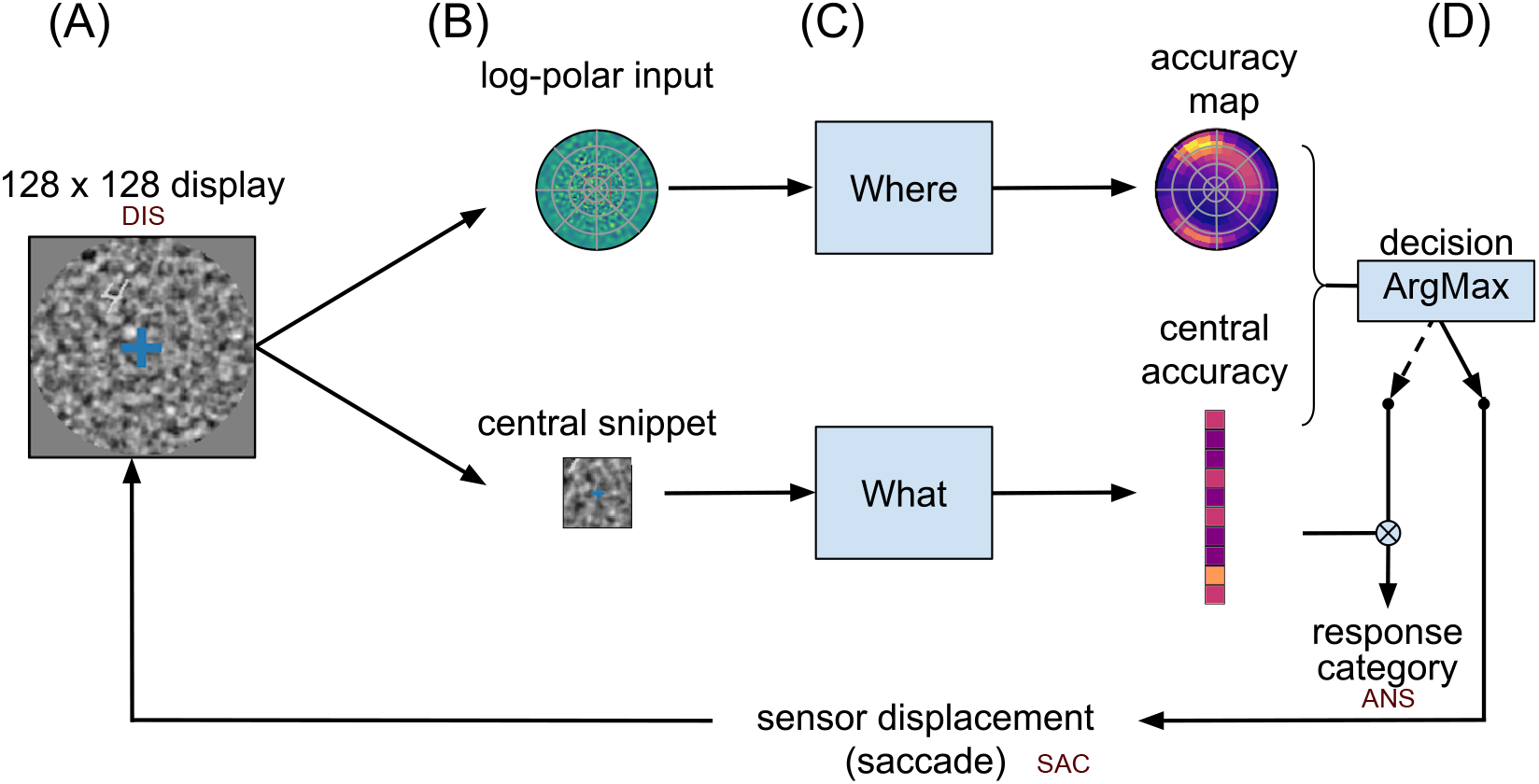
Computational graph. Based on the general anatomy of the visual pathways, we define two streams of information, one stream for identifying a target (“What”?), the other for categorizing it in visual space (“Where”?). **(A)** The visual display is constructed the following way: first a 128 × 128 natural-like background noise is generated, characterized by noise contrast, mean spatial frequency and bandwidth (Sanz-Leon et al., 2012). Then, a sample digit is selected from the MNIST dataset (of size 28 × 28), rectified, multiplied by a contrast factor and overlaid on the background at a random position (see another example in Figure 1-A, `DIS`). Last, a circular mask is put on. **(B)** The visual display is then transformed in 2 sensory inputs: (i) A 28 × 28 central foveal-like snippet is fed to a classification network (“What” pathway). (ii) A log-polar set of oriented visual features is fed to the “Where” pathway. This log-polar input is generated by a bank of filters whose centers are positioned on a log-polar grid and whose radius increases proportionally with the eccentricity. **(C)** The “What” network is implemented using the three-layered LeNet neural network (Lecun et al., 1998). This network outputs a vector predicting the accuracy of detecting the correct digit. In parallel, the “Where” network is implemented by a three-layered neural network consisting of the retinal log-polar input, two hidden layers (fully-connected linear layers combined with a ReLU non-linearity) and a collicular-like accuracy map at the output. This map has a similar log-polar (retinotopic) organization and predicts the accuracy at the hypothetical position of a saccade. Both networks learn to associate the output with the ground truth through back-propagation. **(D)** For a given display, the network provides two accuracy outputs. The two streams converge toward a decision layer that compares the central and the peripheral accuracy which are predicted by both pathways, in order to decide whether to issue a saccadic or a categorical response. If the predicted accuracy in the output of the “What” network is higher than that predicted in the “Where” network, we interrupt the visual search and classify the foveal image using the “What” pathway such as to give the answer (`ANS`). In the other case, the position of maximal activity in the “Where” pathway serves to generate a saccade which shifts the center of gaze. For each saccade realized, the center of vision is displaced and the process is repeated.

In a stationary condition, where the target’s position and identity do not change over time, each saccade thus provides a new viewpoint over the scene, allowing to form a new estimation of the target identity. Following the active inference setup (Friston et al., 2012, Najemnik and Geisler, 2005), we assume that, instead of trying to detect the actual position of the target, the agent tries to maximize the counter-factual benefit in scene understanding that would be gained by any potential saccade. The focus is thus put on action selection metric rather than spatial representation. This means in short estimating how accurate a categorical target classifier will be after moving the eye. In a full setup, predictive action selection means first predicting the future visual field denoted *x′* which is obtained at the center of fixation, and then predicting how good the estimate of the target identity, denoted *y*, i.e. *p*(*y*|*x′*), will be at this location. In practice, predicting a future visual field over all possible saccades is too computationally expensive. Better off instead is to record, for every context *x*, the improvement obtained in recognizing the target after different saccades *a, a′, a″*,…. If *a* is a possible saccade and *x′* the corresponding future visual field, the result of the central categorical classifier over *x′* can either be correct (1) or incorrect (0). If this experiment is repeated many times over many visual scenes, the probability of correctly classifying the future visual field *x′* from *a* is a number between 0 and 1, that reflects the frequency of correct classifications. The putative effect of every saccade can thus be condensed in a single number, the *accuracy*, that quantifies the final benefit of issuing saccade *a* from the current observation *x*. Extended to the full action space *A*, this forms an accuracy map that should monitor the selection of saccades. This accuracy map can be trained by trials and errors, with the final classification success or failure used as a teaching signal. Our main assumption here is that such a *predictive accuracy map* is at the core of a realistic saccade-based vision systems.

From the active inference standpoint, the separation of the scene analysis in two independent tasks relies on a simple “Naïve Bayes” assumption (see Methods). Each processing is assumed to be realized in parallel through different pathways by analogy with the ventral and dorsal pathways in the visual pathways (see Figure 2). A first classifier is thus assigned to process only the pixels found at the center of fixation, while a second one processes the full visual field with a retina-mimetic central log-polar magnification. The first one is called the “What” network, and the second one is the “Where” network (see Figure 7 for details). This combination of a scalar drive with action selection is reminiscent of the actor/critic principle proposed for long time in the reinforcement learning community (Sutton and Barto, 1998). In biology, the ventral and the dorso-lateral division of the striatum have been suggested to implement such an actor-critic separation (Joel et al., 2002, Takahashi et al., 2008). Consistently with those findings, our central accuracy drive and peripheral action selection map can respectively be considered as the “critic” and the “actor” of an accuracy-driven action selection scheme, with foveal identification/disambiguation taken as a “visual reward”.

**Figure 3.**
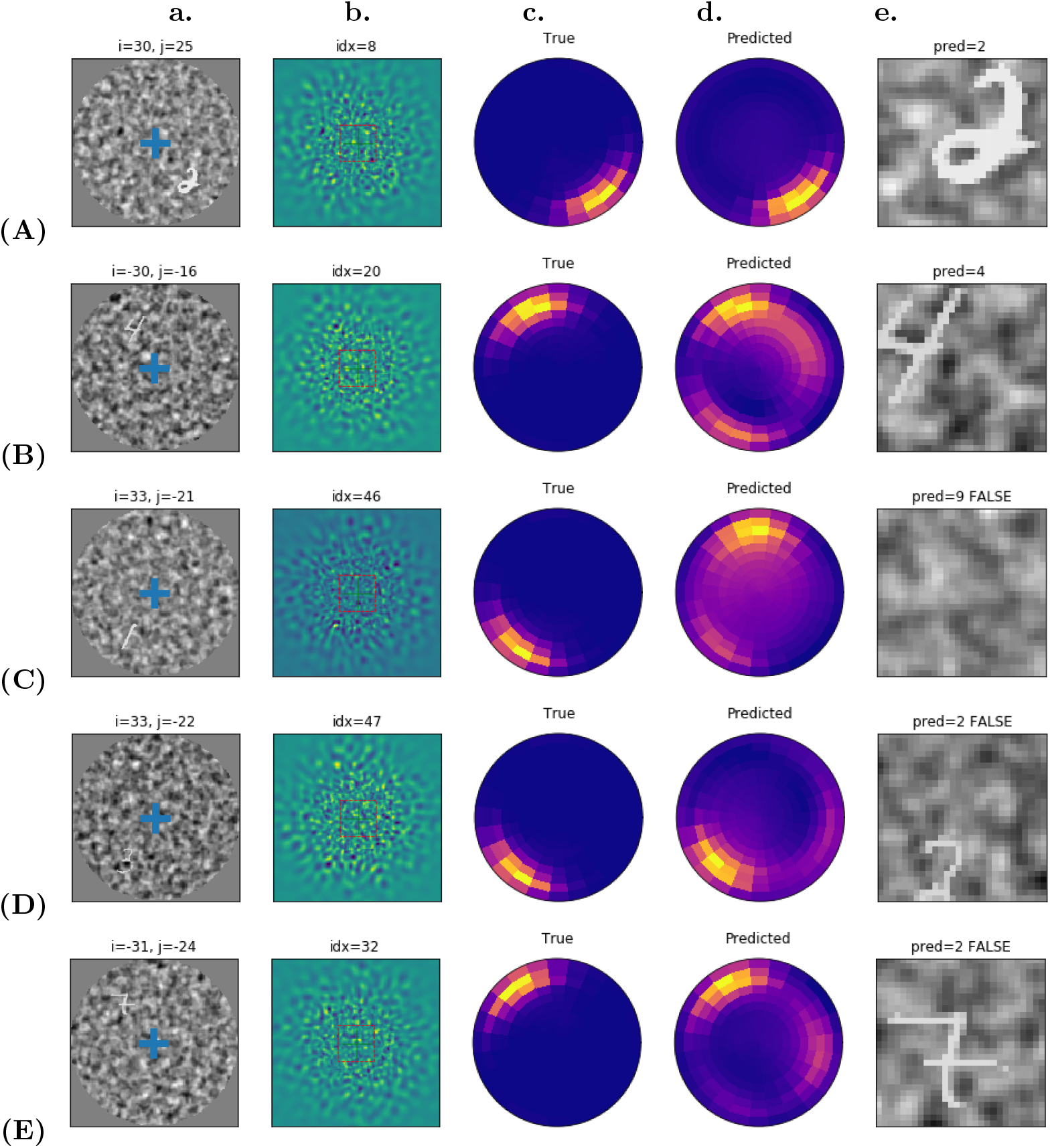
**(A)** – **(E)** Representative samples of active vision after training the “Where” network: **(A)** – **(B)** classification success samples, **(C)** – **(E)** classification failure samples. Digit contrast set to 70%. From left to right: **a.** The initial 128×128 visual display, with blue cross giving the initial center of gaze. The visual input is retinotopically transformed and sent to the multi-layer neural network implementing the “Where” pathway. **b.** Magnified reconstruction of the visual input, as it shows off from the primary visual features through an inverse log-polar transform. **c.-d.** Color-coded radial representation of the output accuracy maps, with dark violet for the lower accuracies, and yellow for the higher accuracies. The network output (“Predicted”) is visually compared with the ground truth (“True”). **e.** The foveal image as the 28 × 28 central snippet extracted from the visual display after doing a saccade, with label prediction and success flag in the title.

**Figure 4.**
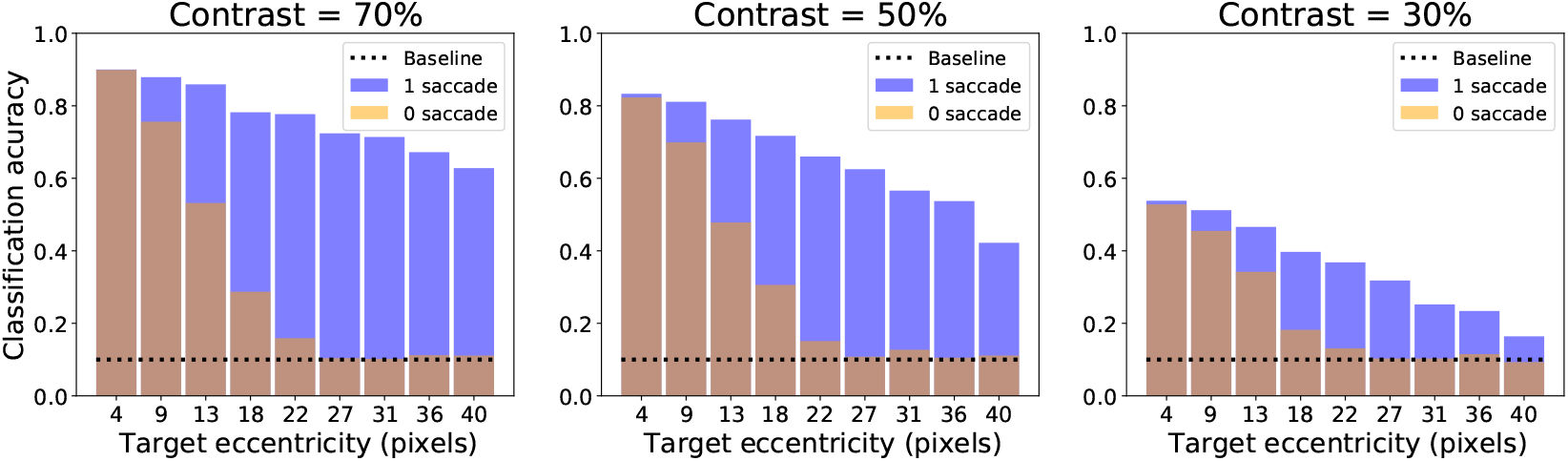
Effect of contrast and target eccentricity. The active vision agent is tested for different target eccentricities (in pixels) and different contrasts. The final classification rate is plotted as transparent orange and blue bars which correspond respectively to the pre-saccadic accuracy from the central classifier (‘0 saccade’) and the post-saccadic accuracy (‘1 saccade’). These are plotted with respect to the target’s eccentricity, and averaged over 1000 trials per eccentricity.

**Figure 5.**
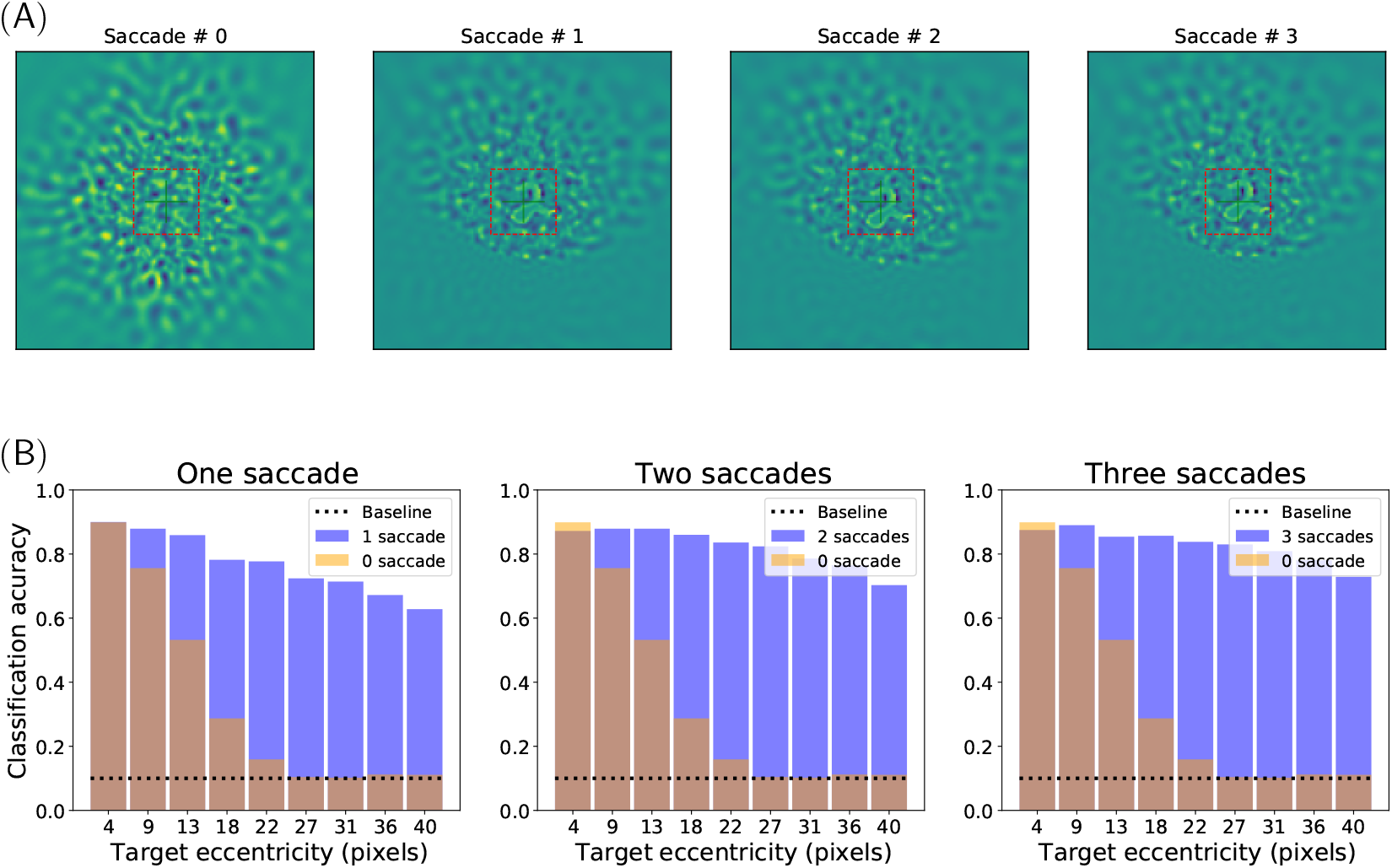
Multiple saccade setup. **(A)** Example of a trial with a sequence of 3 saccades. The subjective visual field is reconstructed from the log-polar visual features, with the red square delineating the 28 × 28 foveal snippet, after 0, 1, 2 and 3 saccades (from left to right). After the first saccade, the accuracy predicted by the “Where” network is higher than that predicted by the “What” network and a corrective saccade is realized to center the target. After this second saccade, the foveal accuracy is higher than that predicted in the periphery and the answer `ANS` is given. **(B)** Average classification accuracies measured for different target eccentricities (in pixels) and a different number of saccades. Target contrast set to 70%. (Transparent) light orange bars: pre-saccadic central accuracy (“0 saccade”) with respect to eccentricity, averaged over 1000 trials per eccentricity. Blue bars: Final classification rate after one, two and three saccades (from left to right, respectively).

**Figure 6.**
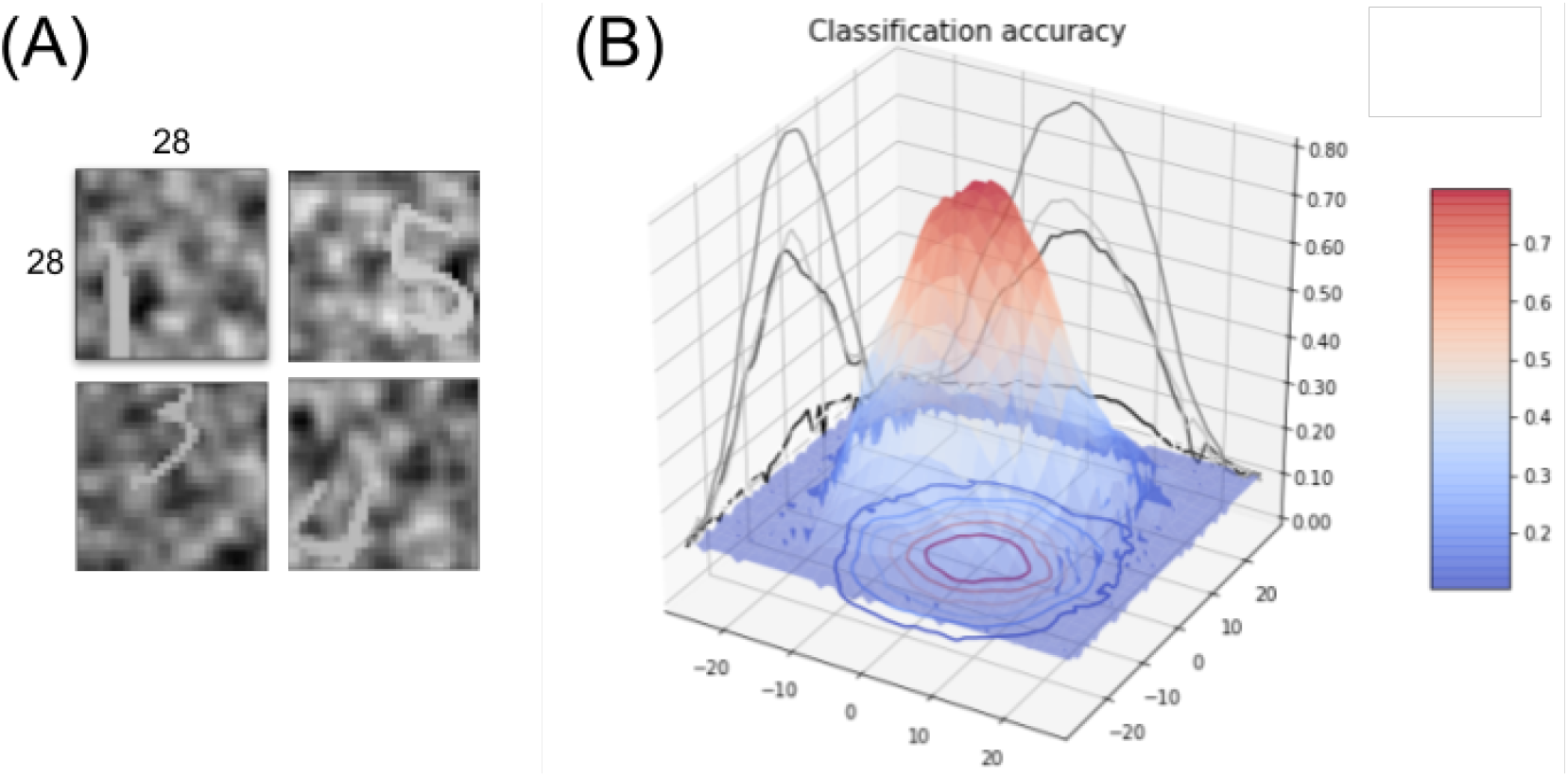
**(A)** Input samples from the “What” training set, with randomly shifted targets using a Gaussian bivariate spatial shift with a standard deviation of 15 pixels. The target contrast is randomly set between 30 % and 70 %. **(B)** 55 × 55 shift-dependent accuracy map, measured for different target eccentricities on the test set after training.

**Figure 7.**
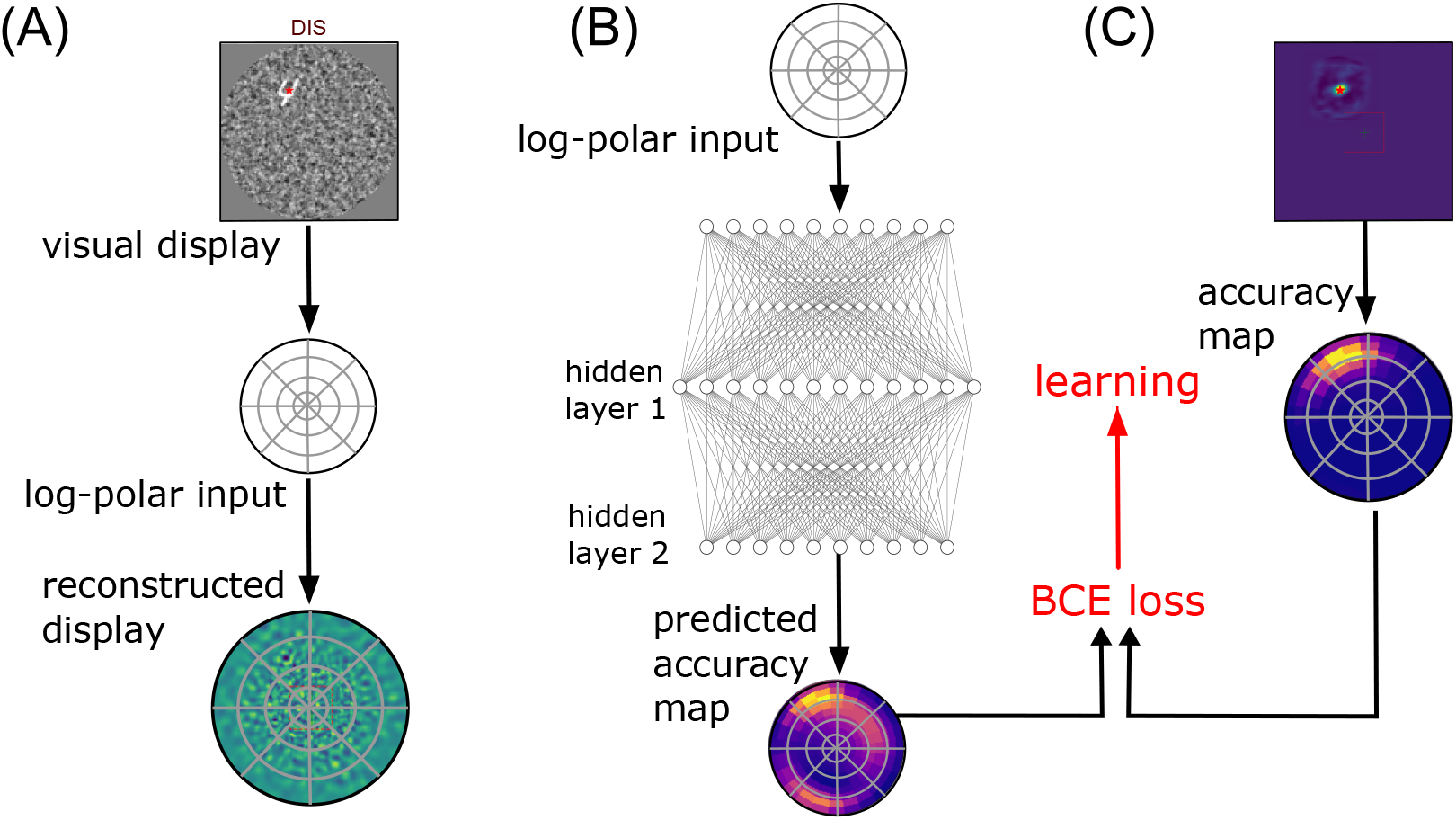
Implementing the “Where” pathway: **(A)** A visual display is transformed by a feature vector which elements compute the similarity of the full image with a bank of oriented filters placed at positions defined by a log-polar grid. This defines a linear transform of the 128 × 128 = 16384 input pixels into 2880 coefficients. It is possible to represent this information in visual space by using the pseudo-inverse of the linear transform (see for instance Figure 1-C). **(B)** The “Where” network consists of two hidden layer composed with a RELU operator transforming the retinal feature vector. A sigmoid operator ensures that this output vector is a distribution of predicted likelihoods in log-polar space. **(C)** Similarly to (A), any full accuracy map computed by shifting the know shift-dependent accuracy map of the “What” pathway (see Figure 6) can be transformed into a distribution in log-polar space, similarly to a collicular representation. As the full accuracy map is itself a distribution, This can be implemented by a linear (matrix) transform. In practice, one can use the inverse of this linear transform to project any collicular representation into the visual space, for instance to predict for the position with maximal accuracy (red cross).

The operations that transform the initial visual data should preserve the initial retinotopic organization, so as to form a final retinotopic accuracy map. Accordingly with the visual data, the retinotopic accuracy map may thus provide more detailed accuracy predictions in the center, and coarser accuracy predictions in the periphery. Finally, each different initial visual field may bring out a different accuracy map, indirectly conveying information about the retinotopic position of the target. A final action selection (motor map) should then overlay the accuracy map through a winner-takes-all mechanism (see Figure 2-D), implementing the saccade selection in a biologically plausible way, as it is thought to be done in the superior colliculus, a brain region responsible for oculomotor control (Sparks and Nelson, 1987). The saccadic motor output showing a similar log-polar compression than the visual input, the saccades should be more precise at short than at long distance and several saccades may be necessary to precisely reach distant targets.

In practice, the “What” and ‘Where” networks are both implemented in `pytorch` (Paszke et al., 2019), and trained with gradient descent over multiple layers. Each network is trained and tested separately. Because the training of the “Where” pathway depends on the accuracy given by the “What” pathway (and not the reverse), we trained the latter first, though a joint learning also yielded similar results. Finally, these are evaluated in a coupled, dynamic vision setup.

## 3 Results

### 3.1 One saccade setup

After training, the “Where” pathway is now capable to predict an accuracy map (see Figure 3), whose maximal argument drives the eye toward a new viewpoint with one saccade. There, a central snippet is extracted, that is processed through the “What” pathway, allowing to predict the digit’s label. Examples of this simple sequence with one saccade are presented in Figure 3, when the digits contrast parameter is set to 70% and the digits eccentricity varies between 0 and 40 pixels. The presented examples correspond to strong eccentricity cases, when the target is hardly visible on the display (Figure 3a), and almost invisible on the reconstructed input (Figure 3b). The radial maps (Figure 3c-d) respectively represent the actual and the predicted accuracy maps. The final focus (foveal area at the location of the selected saccade) is represented in Figure 3e, with cases of classification success (Figure 3A-B) and cases of classification failures (Figure 3C-E). In the case of successful detection (Figure 3A-B), the accuracy prediction is not perfect and the digit is not perfectly centered on the fovea. This “close match” still allows for a correct classification, as the digit’s pixels are fully present on the fovea. The case of Figure 3B and 3C is interesting for it shows two cases of a bimodal prediction, indicating that the network is capable of doing multiple detections in a single pass, that is, *at a glance*. The case of Figure 3C corresponds to a false detection, with the true target detected still, though with a lower intensity. The case of Figure 3D is a “close match” detection that is not precise enough to correctly center the visual target. Some pixels of the digit being invisible on the fovea, the label prediction is mistaken. The last failure case (Figure 3E) corresponds to a correct localization that is harmed by a wrong label prediction, which is due to the “What” classifier inherent error rate.

To test for the robustness of our framework, the same experiment was repeated at different signal-to-noise ratios (SNR) of the input images. Both pathways being interdependent, it is indeed crucial to disentangle the relative effect of both sources of errors in the final accuracy. By manipulating the SNR and the target eccentricity, one can precisely monitor the network detection and recognition capabilities, with a detection task ranging from “easy” (small shift, strong contrast) to “highly difficult” (large shift, low contrast). The digit recognition capability is systematically evaluated in Figure 4 for different eccentricities and different SNRs. We test the final accuracy of the system for three target contrasts conditions ranging from 30% to 70% of the maximal contrast, and 10 different eccentricities ranging from 4 to 40 pixels. It is averaged over 1,000 trials both on the initial central snippet and the final central snippet (that is, at the landing of the saccade). The (transparent) orange bars provide the initial classification rate (without saccade) and the blue bars provide the final classification rate (after saccade) – see Figure 4. As expected, the accuracy decreases in both cases with the eccentricity, for the targets become less and less visible in the periphery. The decrease is rapid in the pre-saccadic case: the accuracy drops to the baseline level for a target distance of approximately 20 pixels from the center of gaze. The post-saccadic accuracy has a much wider range, with a slow decrease up to the border of the visual display (40 pixels away from the center). When varying the target contrast, the pre-saccadic accuracy profile is scaled by the reference accuracy (obtained with a central target), whose values are approximately 92%, 82% and 53% for contrasts of 70, 50 and 30%. The post-saccadic accuracy profile undergoes a similar scaling at the different contrast values, indicating the critical dependence of the global setup to the central processing reliability.

The high contrast case (see Figure 4) provides the greatest difference between the two profiles, with an accuracy approaching 90% at the center and 60% at the periphery. This allows to recognize digits after one saccade in a majority of cases, up to the border of the image, from a very scarce peripheral information. With decreasing target contrast, a general decrease of the accuracy is observed, both at the center and at the periphery, with about 10% decrease with a contrast of 50%, and 40% decrease with a contrast of 30%. In addition, the proportion of false detections also increases as contrast decreases. At 40 pixels away from the center, the false detection rate is approximately 40% for a contrast of 0.7, 60% for a contrast of 0.5 and 80% for a contrast of 0.3 (with a recognition close to the baseline at the periphery in that case). The difference between the initial and the final accuracies is maximal for eccentricities ranging from 15 to 30 pixels. This optimal range reflects a proportion of the visual field around the fovea where the target detection is possible, but not its identification. The visual agent knows *where* the target is, without exactly knowing *what* it is.

### 3.2 Analysis

This full covering of the 128×128 image range is done at a much lesser cost than what would be done by a systematic image scan, as in classic computer vision. Taking *n* the number of pixels in the original image (in our case *n* = 128 × 128 = 16384), our log-polar encoding provides *O*(log *n*) log-polar visual features by construction. The total visual data processed is the addition of the *C* pixels processed at the fovea and the *O*(log *n*) log-polar visual features processed at the periphery. The total processing cost is thus *O*(*C* + log *n*). Taking *C* as a constant, the total processing cost can be said *O*(log *n*) (for constant processing times do not change the order). In the case of multiple saccades (see next section), the total cost is *O*(*k* × (*C* + log *n*)) with *k* the number of saccades. If the number of saccades *k* is bounded by a constant *K*, this allows to estimate the processing cost as *O*(*K* × (*C* + log *n*)) in the worst case, that also resumes to *O*(log *n*). This is to be contrasted, for instance, with the linear cost obtained with a full convolutional scan with a window of size *C* and a stride of 1, that is precisely *O*(*C* × *n*). Various optimizations can of course be considered, of which the well-known max-pooling principle used in deep learning, but anyway image processing (without compression loss) is generally considered as linear in the size of the visual data processed (Strengert et al., 2006)

Our sub-linear processing time thus justifies a strategy that may have been chosen in a variety of natural vision systems. The compromise between the urgency to detect and the need to be accurate may justify the different balances which may exist in different species. In particular, this may justify the differences observed between preys (with a less sparse cone density at the periphery) and predators (with a tendency toward denser foveal regions).

### 3.3 Multiple saccades setup

In our simulation results, the post-saccadic accuracy is found to overtake the pre-saccadic accuracy *except* when the target is initially close to the center of gaze. When closely inspecting the 1-10 pixels eccentricity range in our first experiment (not shown), a decision frontier between a positive and a negative information gain is found at 2-3 pixels away from the center. Inside that range, no additional saccade is expected to be produced, and a categorical response should be given instead. It is crucial here to understand that this empirical accuracy difference can be predicted, by construction, as the difference of the maximal outputs of the “Where” and the “What” pathways. This difference-of-accuracies prediction can serve as a decision criterion before actuating the saccade, like a GO/NOGO signal. It is moreover interpretable as an approximation of the information gain provided by the “Where” pathway, with the true label log-posterior seen as a sample of the posterior entropy (see Equation 1 in section 5.5).

After the first saccade, while the decision criterion is not attained, additional saccades may be pursued in order to search for a better centering. In the case of a false detection for instance, the central accuracy estimate should be close to the baseline, and may allow to “explain away” the current center of gaze and its neighborhood, encouraging to actuate long-range saccades toward less salient peripheral positions, making it possible to escape from initial prediction errors. This incitement to select a saccade “away” from the central position is reminiscent of a well-known phenomenon in vision known as the “inhibition of return” (Itti and Koch, 2001). Combining accuracy predictions from each pathway may thus allow to refine saccades selection in a way that complies with the sequential processing observed in biological vision. Note that extended to a multi-target case, the Information Gain maximization principle still holds as a general measure of the improvement of scene understanding through multiple saccades. It is uncertain however wether biologically realistic implementations would be possible in that case. In particular, we predict that such a mechanism is dependent on the class of inputs, and would be for instance different when searching for faces as compared to digits.

Some of the most peripheral targets are thus difficult to detect in just one saccade, resulting in degraded performances at the periphery (see Figure 4). Even when correctly detected, our log-polar action maps also precludes precise centering. As a consequence, peripheral targets are generally poorly centered after the first saccade, as shown for instance in Figure 3-D, resulting in classification errors. The possibility to perform a sequential search using more saccades is thus crucial to allow for a better recognition. Results on multi-saccades visual search results are presented in Figure 5. An example of a trial with a sequence of 3 saccades is shown in Figure 5-A. A hardly visible peripheral target (digit) is first approximately shifted to the foveal zone thanks to the first saccade. Then, a new retinal input centered at the new point of fixation is computed, such that it generates a novel predicted accuracy map. The second saccade allows to improve the target centering. As the predicted foveal accuracy given by the “What” network is higher than the peripheral one given by the “Where” network, a third saccade would not improve the centering: The stopping criteria is met. In practice, 1 or 2 saccades were sufficient in most trials to reach the actual target. Another behavior was also observed for some “bad start” which exhibited a false localization (as in Figure 3-C for instance), when the target is shifted away in the opposite direction and the agent can not recover from its initial error. From Figure 5-B, this case can be estimated at about 15% of the cases for the most peripheral targets.

Overall, as shown in Figure 5-B, the corrective saccades implemented in this multiple saccade setup provide a significant improvement in the classification accuracy. Except at the center, the accuracy increases by about 10% both for the mid-range and the most peripheral eccentricities. Most of the improvement however is provided by the first corrective saccade. The second corrective saccade only shows a barely significant improvement of about 2% which is only visible at the periphery. The following saccades would mostly implement target tracking, without providing additional accuracy gain. A 3-saccades setup finally allows a wide covering of the visual field, providing a close to central recognition rate at all eccentricities, with the residual peripheral error putatively corresponding to the “bad start” target misses cases.

## 4 Discussion

### 4.1 Summary

In summary, we have proposed a visuomotor action-selection model that implements a focal accuracy-seeking policy across the image. Our main modeling assumption here is an *accuracy-driven* monitoring of action, stating in short that the ventral classification accuracy drives the dorsal selection in building an extra-foveal accuracy map. The comparison of both accuracies amounts either to select a saccade or to keep the eye focused at the center, so as to identify the target. The predicted accuracy map has, in our case, the role of a value-based action selection map, as it is the case in model-free reinforcement learning. However, it also owns a probabilistic interpretation, making it possible to combine concurrent accuracy predictions, such as the ones done through the “What” and the “Where” pathways. This allows in particular to explain more elaborate aspects of the whole decision making processes, such as the inhibition of return (Itti and Koch, 2001), without further specific heuristic.

Moreover, one crucial aspect highlighted by our model is the importance of centering objects in recognition. Despite the robust translation invariance observed on the “What” pathway, a small tolerance radius of about 4 pixels around the target’s center needs to be respected to maximize the classification accuracy. The translation invariance is in our case an effect of the max-pooling operations in the convolutional layers, build-in at the core of the “What” layer. This relates to the idea of finding an absolute referential for an object, for which the recognition is easier. If the center of fixation is fixed, the log-polar encoding of an object has the notable properties to map object rotations and scalings toward translations in the radial and angular directions of the visual domain (Javier Traver and Bernardino, 2010). Extensions to scale and rotation invariance would in principle be feasible through central log polar encoding, with little additional computational cost. This prospect is left for future work.

### 4.2 Comparison with other models

A lot of computer models found in the literature reflect to some degree the foveal/sequential visual processing principles developed here. Since the question of a normative and quantitative comparison with them is important, no specific or unified dataset is proposed at present to address this specific case. Every model found uses a different retinal encoding, different computing methodologies and different training datasets. We thus provide here a qualitative comparison with the more prominent computer-based focal vision models proposed in the literature.

First, active vision is of course an important topic in mainstream computer vision. In the case of image classification, it is considered as a way to improve object recognition by progressively increasing the definition over identified regions of interest, referred as “recurrent attention” (Fu et al., 2017, Mnih et al., 2014). Standing on a similar mathematical background, recurrent attention is however at odd with the functioning of biological systems, with a mere distant analogy with the retinal principles of foveal-surround visual definition. Phenomenological models, such as the one proposed in Najemnik and Geisler’s seminal paper (Najemnik and Geisler, 2005), rely on a rough simplification, with foveal center-surround acuity modeled as a response curve. Despite providing a bio-realistic account of sequential visual search, the model owns no foveal image processing implementation. Stemming on Najemnik and Geisler’s principles, a trainable center-surround processing system was proposed in (Butko and Movellan, 2010), with a sequential scan of an image in a face-detection task. However, the visual search task relies there on a systematic scan over a dynamically-blurred image, with all the visual processing delegated to standard feature detectors.

In contrast, the Akbas and Eckstein model (“foveated object detector” (Akbas and Eckstein, 2017)) uses an explicit bio-inspired log-polar encoding for the peripheral processing, with trainable local features. With a focus put on the processing effectiveness provided by this specific compression, the model approaches the performance of state-of-the-art linear feature detectors, with multi-scale template matching (bounding box approach). However the use of a local/linear template matching processing makes here again the analogy with the brain oversimplistic.

Denil et al’s paper (Denil et al., 2012) is probably the one that shows the closest correspondence with our setup. It owns an identity pathway and a control pathway, in a What/Where fashion, just as ours. Interestingly, only the “What” pathway is neurally implemented using a random foveal/multi-fixation scan within the fixation zone. The “Where” pathway, in contrast, mainly implements object tracking, using particle filtering with a separately learned generative process. The direction of gaze is here chosen so as to minimize the target’s position, speed and scale uncertainty, using the variance of the future beliefs as an uncertainty metric. The control part is thus much similar to a dynamic ROI tracking algorithm, with no direct correspondence with foveal visual search, or with the capability to recognize the target

### 4.3 Perspectives

We have thus provided here a proof of concept that a log-polar retinotopy can efficiently serve object detection and identification over wide visual displays. Despite its simplicity, the model used to generate our visual display allowed to assess the effectiveness and robustness of our learning scheme, that should be extended in the future to more complex displays and more realistic closed-loop setups. In particular, the restricted 28 × 28 input used for the foveal processing is a mere placeholder, that should be replaced by more elaborate computer vision frameworks, such as Inception (Szegedy et al., 2015) or VGG-19 (Simonyan and Zisserman, 2014), that can handle more ecological natural image classification setups.

The main advantage of our peripheral image processing is its cost-efficiency. Our full log-polar processing pathway consistently conserves the high compression rate performed by retina and V1 encoding up to the action selection level. The organization of both the visual filters and the action maps in concentric log-polar elements, with radially exponentially growing spatial covering, can thus serve as a baseline for a future sub-linear (logarithmic) complexity for visual search in computer vision. Our work thus illustrates one of the main advantages of using a focal/sequential visual processing framework, that is providing a way to process large images with a sub-linear processing cost. This may allow to detect an object in large visual environments, which should be particularly beneficial when the computing resources are under constraint, such as for drones or mobile robots.

If the methodology and principles developed here are clearly intended to deal with real images, an important contribution of the paper is providing principles that justify the separation between a ventral and a dorsal stream in the early visual pathways. If some forms of “dual pathway models” have been proposed in the past (through separating the central and the peripheral processing, like in (Denil et al., 2012), and also in one instance of the (Akbas and Eckstein, 2017) model, their guiding principles stem on computing efficacy rather than biological fidelity. We thus think that our principled ventral/dorsal concurrent processing, rooted on dorsal accuracy map predictions, is both important and novel.

Finally, our approach relies on a strong idealization, assuming the presence of a unique target. This is well adapted to a fast changing visual scene as is demonstrated by our ability to perform as fast as 5 saccades per second to detect faces in a cluttered environment (Martin et al., 2018). However, some visual scenes—such as when looking at a painting in a museum—allow for a longer inspection of its details. The presence of many targets in a scene should be addressed, which amounts to sequentially select targets, in combination with implementing a more elaborate inhibition of return mechanism to account for the trace of the performed saccades. This would generate more realistic visual scan-paths over images. Actual visual scan-paths over images could also be used to provide priors over action selection maps that should improve realism. Identified regions of interest may then be compared with the baseline bottom-up approaches, such as the low-level feature-based saliency maps (Itti and Koch, 2001). Maximizing the Information Gain over multiple targets needs to be envisioned with a more refined probabilistic framework extending previous models (Friston et al., 2012), which would include phenomena such as mutual exclusion over overt and covert targets. Next, scan-paths could be generated on actual dynamical scenes, and this despite existing oculomotor delays (Perrinet et al., 2014), extending such a framework in the temporal domain. How the brain may combine and integrate these various probabilities dynamically is still an open question, that amounts to the fundamental binding problem.

## 5 Methods

### 5.1 Image generation

We first define here the generative model for input display images as shown first in Figure 1-A (`DIS`) and as implemented in Figure 2-A. Following a common hypothesis regarding active vision, visual scenes consist of a single target embedded in a large image with a cluttered background.

#### Targets

We use the MNIST dataset of handwritten digits introduced by (Lecun et al., 1998): Samples are drawn from the dataset of 60000 grayscale, 28 × 28 pixels images. There are 10 categories (from “zero” to “nine”). These are separated between a training and a validation set (see below the description of the “Where” network). Note that to simplify the task, there is one and only one target per image.

#### Full-scale images

We call “full-scale images”, input images which correspond to a discretized, rectangular sampling (pixels) of the visual field, such that which are usually used in computer vision. These input images are set to a size of 128 × 128 pixels in which we embed the target. Each target location is drawn at random in this large image. To enforce isotropic generation (at any direction from the fixation point), a centered circular mask covering the image (of radius 64 pixels) is defined. Also, the target’s location is such that the embedded sample fits entirely into that circular mask.

#### Background noise setting

To implement a realistic background noise, we generate synthetic textures (Sanz-Leon et al., 2012) using a bi-dimensional random process. The texture is designed to fit with the statistics of natural images. We chose an isotropic setting where textures are characterized by solely two parameters, one controlling the median spatial frequency of the noise, the other controlling the bandwidth around the central frequency. Equivalently, this can be considered as the band-pass filtering of a random white noise image. The spatial frequency is set at 0.1 pixel^-1^ to fit that of the original digits. This specific spatial frequency occasionally allows to generate some “phantom” digit shapes in the background. Finally, these images are rectified to have a normalized contrast.

#### Mixing the signal and the noise

Finally, both the noise and the target image are merged into a single image. Two different strategies are used. A first strategy emulates a transparent association, with an average luminance computed at each pixel, while a second strategy emulates an opaque association, choosing for each pixel the maximal value. The quantitative difference was tested in simulations, but proved to have a marginal importance and results shown here are the result of the opaque association.

### 5.2 Active inference and the Naïve Bayes assumption

Saccade selection in visual processing can be captured by a statistical framework called a partially observed Markov Decision Process (POMDP) (Butko and Movellan, 2010, Friston et al., 2012, Najemnik and Geisler, 2005), where the cause of a visual scene is made up from the couple of independent random variables of the viewpoint and of the scene elements (here a digit). For instance, changing the viewpoint will lead to a different scene rendering. A generative model tells how the visual field should look knowing the scene elements and a certain viewpoint. In general, active inference assumes a hidden external state e, which is known indirectly through its effects on the sensor. The external state corresponds to the physical environment. Here the external state is assumed to split in two (independent) components, namely *e* = (*u, y*) with *u* the interoceptive body posture (in our case the gaze orientation, or “viewpoint”) and *y* the object shape (or object identity). The visual field *x* is the state of the sensors, that is, a partial view of the visual scene, measured through the generative process: *x* ~ *p*(*X*|*e*).

Using Bayes rule, one may then infer the scene elements from the current viewpoint (model inversion). The real physical state *e* being hidden, a parametric model *θ* is assumed to allow for an estimate of the cause of the current visual field through model inversion thanks to Bayes formula: *p*(*E*|*x*) ∝ *p*(*x*|*E*; *θ*). It is also assumed that a set of motor commands *A* = {…, *a*,…} (here saccades) may control the body posture (gaze orientation), but not the object’s identity, so that *y* is independent of *a*. Actuating a command *a* changes the viewpoint to *u′*, which feeds the system with a new visual sample *x′* ~ *p*(*X* |*u′, y*). The more viewpoints you have, the more certain you are about the object identity through a chain rule sequential evidence accumulation.

In an optimal search setup however (Najemnik and Geisler, 2005), you need to choose the next viewpoint that will help you to disambiguate *at best* the scene. In a predictive setup, the consequence of every saccade should be analyzed through model inversion *over the future observations*, that is, predicting the effect of every action to choose the one that may optimize future inferences. The benefit of each action should be quantified through a certain metric (future accuracy, future posterior entropy, future variational free energy, …), that depend on the current inference *p*(*U, Y*|*x*). The saccade *a* that is selected thus provides a new visual sample from the scene statistics. If well chosen, it should improve the understanding of the scene (here the target position and category). However, estimating in advance the effect of every action over the range of every possible object shapes and body postures is combinatorially hard, even in simplified setups, and thus infeasible in practice.

The predictive approach necessitates in practice to restrain the generative model in order to reduce the range of possible combinations. One such restriction, known as the “Naïve Bayes” assumption, considers the independence of the factors that are the cause of the sensory view. The independence hypothesis allows considering the viewpoint *u* and the category *y* being independently inferred from the current visual field, i.e *p*(*U, Y*|*x*) = *p*(*U*|*x*)*p*(*Y*|*x*). This property is strictly true in our setting and is very generic in vision for simple classes (such as digits) and simple displays (but see (Võ and Wolfe, 2012) for more complex visual scene grammars).

### 5.3 Foveal vision and the “What” pathway

At the core of the vision system is the identification module, i.e. the “What” pathway (see Figure 2). It consists of a classic convolutional classifier for which we will show some translation invariance in the form of a shift-dependent accuracy map. Importantly, it can quantify its own classification uncertainty, that may allow comparisons with the output of the “Where” pathway.

The foveal input is defined as the 28 × 28 grayscale image cropped at the center of gaze (see dashed red box in Figure 1-C). This image is passed unmodified to the agent’s visual categorical pathway (the “What” pathway), that is realized by a convolutional neural network, here the well-known “LeNet” classifier (Lecun et al., 1998). The network structure that processes the input to identify the target category is made of three convolution layers interleaved with max-pooling layers, followed by two fully-connected layers as provided (and unmodified) by its Pytorch library implementation (Paszke et al., 2019). Each intermediate layer’s output is rectified and the network output uses a sigmoid operator to predict the likelihood of detecting each of the 10 digits. The index of one of the 10 output neuron with maximum probability provides the image category. It is first trained over the (centered) MNIST dataset after approx 20 training epochs. This strategy achieves an average 98.7% accuracy in the center on the validation dataset (Lecun et al., 1998).

To achieve an even more generic “What” pathway, a specific dataset is constructed to train the network. It is made of randomly shifted digits overlayed over a randomly generated noisy background, as defined above. Both the shift and the contrast relative to the background noise make the task more difficult than the original MNIST categorization. The relative contrast of the digit is randomly set between 30 % and 70 % of the maximal contrast. The network is trained incrementally by progressively increasing the shift variability (of a bivariate central gaussian) and by increasing the standard deviation from 0 to 15 (with a maximal shift set at 27 pixels). The network is trained on a total of 75 epochs, with 60000 examples generated at each epoch from the original MNIST training set, using the cross-entropy loss. The shifts and backgrounds are re-generated at each epoch. The shifts’ standard deviation increases of one unit every 5 epochs such that at the end of the training, many digits fall outside the center of the fovea, so that many examples are close to impossible to categorize, either because of a low contrast or a too large eccentricity. At the end of the training process, the average accuracy is thus of 34% and a maximum accuracy 91% at the center.

After training, this shift-dependent accuracy map is validated by systematically testing the network accuracy on every horizontal and vertical shifts, each on a set of 1000 cluttered target samples generated from the MNIST test set and within the range of ±27 pixels (see Figure 6). This forms a 55 × 55 accuracy map showing higher accuracy at the center, and a slow decreasing accuracy with target eccentricity (with an accuracy plateau over 70% showing a relative shift invariance on around 7 pixels eccentricity radius). This shift invariance is a known effect of convolutional computation. Note that the categorization task is here harder by construction and the accuracy that is obtained here is lower (with a central recognition rate of around 80%). The accuracy sharply drops for eccentricities greater than 10 pixels, reaching the baseline 10% chance level at shift amplitudes at around 20 pixels.

### 5.4 “Where” pathway: Transforming log-polar feature vectors to log-polar action maps

Here, we assume the “Where” implements the following action selection: where to look next in order to reduce the uncertainty about the target identity? The “Where” pathway is thus devoted to choosing the next saccade by predicting the location of the target in the (log-polar) visual field. This implies moving the eye such as to increase the “What” categorization accuracy. For a given visual field, each possible future saccade has an expected accuracy,that can be trained from the “What” pathway output. To accelerate the training, we use an equivalent strategy by training the network on a translated accuracy map (see below for details). The output is thus an accuracy map, that tells for each possible visuomotor displacement the value of the future accuracy.

#### Primary visual representation: log-polar orientation filters

In order to reduce the processing cost, and in accordance with observations (Connolly and Van Essen, 1984, Sparks and Nelson, 1987), a similar log-polar compression pattern is assumed to be conserved from the retina up to the visuo-motor layers. The non-uniform sampling of the visual space is adequately modeled as a log-polar conformal mapping, as it provides a good fit with observations in mammals (Javier Traver and Bernardino, 2010) and which has a long history in computer vision and robotics. Both the visual features and the output accuracy map are to be expressed in retinal coordinates. On the visual side, local visual features are extracted as oriented edges as a combination of the retinotopic transform with primary visual cortex filters (Fischer et al., 2007), see Figure 7-A. The centers of these first and second order orientation filters are radially organized around the center of fixation, with small and tightened receptive fields at the center and more large and scarce receptive fields at the periphery. The size of the filters increases proportionally to the eccentricity. The filters are organized in 10 spatial eccentricity scales (respectively placed at around 2, 3, 4.5, 6.5, 9, 13, 18, 26, 36.5, and 51.3 pixels from the center) and 24 different azimuth angles allowing them to cover most of the original 128 × 128 image. At each of these positions, 6 different edge orientations and 2 different phases (symmetric and anti-symmetric) are computed. This finally implements a (fixed) bank of linear filters which models the receptive fields of the primary visual cortex.

To ensure the balance of the coefficients across scales, the images are first whitened and then linearly transformed into a retinal input as a feature vector ***x***. The length of this vector is 2880, such that the retinal filter compresses the original image by about 83%, with high spatial frequencies preserved at the center and only low spatial frequencies conserved at the periphery. In practice, the bank of filters is pre-computed and placed into a matrix for a rapid transformation of input batches into feature vectors. This matrix transformation allows also the evaluation of a reconstructed visual image given a retinal activity vector thanks to a pseudo-inverse of the forward transform matrix. In summary, the full-sized images are transformed into a primary visual feature vector which is fed to the “Where” pathway.

#### Visuo-motor representation: “Collicular” accuracy maps

The output of the “Where” pathway is defined as an *accuracy map* representing the recognition probability after moving the eye, independently of its identity. Like the primary visual map, this target accuracy map is also organized radially in a log-polar fashion, making the target position estimate more precise at the center and fuzzier at the periphery. This modeling choice is reminiscent of the approximate log-polar organization of the superior colliculus (SC) motor map (Sparks and Nelson, 1987). To ensure that this output is a distribution function, we use a sigmoid operator at the ouput of the “Where” network. In ecological conditions, this accuracy map should be trained by sampling, i.e. by “trial and error”, using the actual recognition accuracy (after the saccade) to grade the action selection. For instance, we could use corrective saccades to compute (a posteriori) the probability of a correct localization. In a computer simulation however, this induces a combinatorial explosion which makes the calculation not amenable.

In practice, as we designed the generative model for the visual display, the position of the target (which is hidden to the agent) is known. Combining this translational shift and the shift-dependent accuracy map of the “What” classifier (Figure 6-B), the full accuracy map at each pixel can be thus predicted for each visual sample under an ergodic assumption by shifting the central accuracy map on the true position of the target (see Figure 7-C). Such a computational shortcut is allowed by the independence of the categorical performance with position. This full accuracy map is a probability distribution function defined on the rectangular grid of the visual display. We project this distribution on a log-polar grid to provide the expected accuracy of each hypothetical saccade in a retinotopic space similar to a collicular map. In practice, we used Gaussian kernels defined in the log-polar space as a proxy to quantify the projection from the metric space to the retinotopic space. This generates a filter bank at 10 spatial eccentricies and 24 different azimuth angles, i.e. 240 output filters. To ensure keeping a distribution function, each filter is normalized such that the value at each log-polar position is the average of the values which are integrated in visual space. Applied to the full sized ground truth accuracy map computed in metric space, this gives an accuracy map at different location of a retinotopic motor space.

#### Classifier training

The “Where” pathway is a function transforming an input retinal feature vector ***x*** into an output log-polar retinotopic vector ***a*** representing for each area of the log-polar visual field a prediction of the accuracy probability. Following the active inference framework, the network is trained to predict the likelihood ***a**_i_* at position *i* knowing the retinal input **x** by comparing it to the known ground truth distribution computed over the motor map. The loss function that comes naturally is the Binary Cross-Entropy. At each individual position i, this loss corresponds to the negative term of Kullback-Leibler divergence for a binomial random variable ***a**_i_* given by the predicted map and the ground truth (see Figure 7-B). The total loss is the average over all positions *i*. This scalar measures the distance between both distributions, it is always positive and null if and only if they are equal.

The parametric neural network consists of a primary visual input layer, followed by two fully connected hidden layers of size 1000 with rectified linear activation, and a final output layer with a sigmoid nonlinearity to ensure that the output is compatible with a likelihood function (see Figure 7-B). An improvement in convergence speed was obtained by using batch normalization. The network is trained on 60 epochs of 60000 samples, with a learning rate equal to 10^-4^ and the Adam optimizer (Kingma and Ba, 2014) with standard momentum parameters. One full training takes about 1 hour on a laptop. The code is written in Python (version 3.7.6) with pyTorch library (Paszke et al., 2019) (version 1.1.0). The full scripts for reproducing the figures and explore the results to the full range of parameters is available at https://github.com/laurentperrinet/WhereIsMyMNIST.

#### Quantitative role of parameters

In addition, we controlled that the training results are robust to changes in an individual experimental or network parameters from the default parameters (see Figure 8). From the scan of each of these parameters, the following observations were remarkable. First we verified that accuracy decreased when `noise` increased and while the bandwidth of the noise imported weakly, the spatial frequency of the noise was an important factor. In particular, final accuracy was worst for a clutter spatial frequency of ≈ 0.07, that is when the characteristic textures elements were close to the characteristic size of the objects. Second, we saw that the dimension of the “Where” network was optimal for a dimensionality similar to that of the input but that this mattered weakly. The dimensionality of the log-polar map is more important. The analysis proved that an optimal accuracy was achieved when using a number of 24 azimuthal directions. Indeed, a finer log-polar grid requires more epochs to converge and may result in an over-fitting phenomenon hindering the final accuracy. Such fine tuning of parameters may prove to be important in practical applications and to optimize the compromise between accuracy and compression.

**Figure 8.**
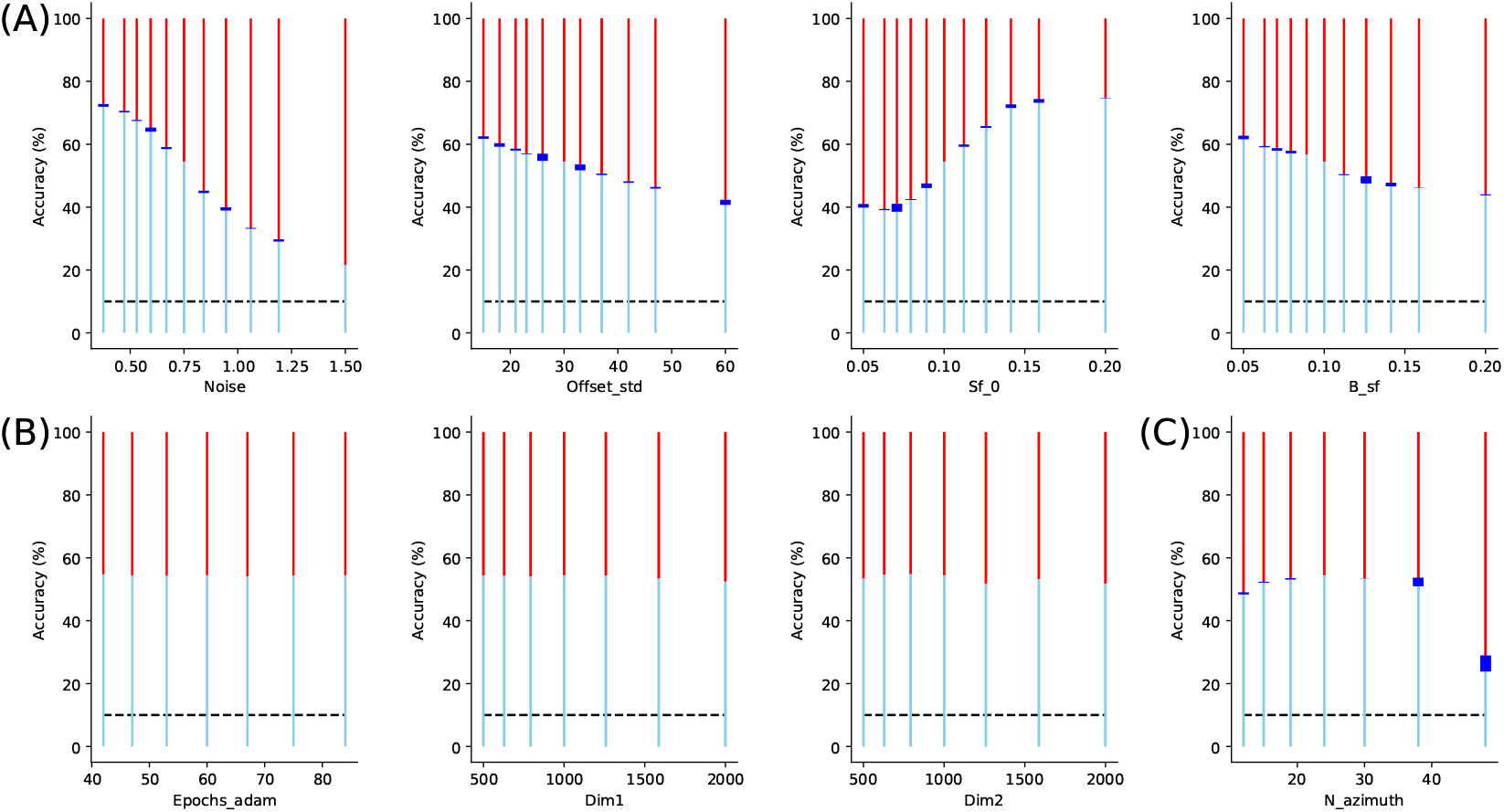
Quantitative role of parameters: We tested all parameters of the presented model, from that controlling the architecture of image generation, to the parameters of the neural network implementing the “Where” pathway (including meta-parameters of the learning paradigm). We show here the results which show the most significative impact on average accuracy. The accuracy is given by a blue line, the red line giving the rate of errors. The black dashed line gives the chance level (10%), while the blue box gives the 99% confidence interval as estimated over 8 repetitions of the learning. **(A)** First, we tested some properties of the input, respectively from left to right: noise level (`Noise`), standard deviation of the distance of the target with respect to the fixation (`Offset_std`), mean spatial frequency of clutter (`Sf_0`) and bandwidth (`B_sf`) of the clutter noise. This shows that average accuracy evolves with noise (see also Figure 4 for an evolution as a function of eccentricity), but also to the characteristics of the noise clutter. In particular, there is a drop in accuracy whenever noise is of similar wavelength as digits, but which becomes less pronounced as the bandwidth increases. **(B)** Finally, we scanned parameters of the Deep Learning neural network. We observed that accuracy quickly converged after approximately 25 epochs (`Epochs_adam`). We then tested different values for the dimension of respectively the first (`Dim1`) and second (`Dim2`) hidden layers, showing weak changes in accuracy. **(C)** The accuracy also changes with the architecture of the foveated input as shown here by changing the number `N_azimuth` of azimuth directions which are sampled in visual space. This shows a compromise between a rough azimuth representation and a large precision, which necessitates a longer training phase, such that the optimal number is around 24 azimuth directions.

### 5.5 Concurrent action selection

Finally, when both pathways are assumed to work in parallel, each one may be used concurrently to choose the most appropriate action. Two concurrent accuracies are indeed predicted through separate processing pathways, namely the central pixels recognition accuracy through the “What” pathway, and the log-polar accuracy map through the “Where” pathway. The central accuracy may thus be compared with the maximal accuracy as predicted by the “Where” pathway.

From the information theory standpoint, each saccade comes with fresh visual information about the visual scene that can be quantified by a conditional *information gain*, namely:

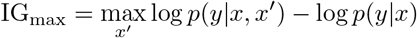

with the left term representing the future accuracy (after the saccade is realized) and the right term representing the current accuracy as it is obtained from the “What” pathway. Estimating the joint conditional dependence in the first term being once again out of reach for computational reasons, the following approximative estimate is used instead:

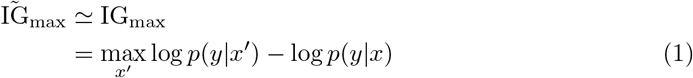

that is a simple difference between the log accuracy after the saccade minus the log accuracy before the saccade. To provide a reliable estimate, the information gain may be averaged over many saccades and many target eccentricities (so that the information gain may be close to zero when the target eccentricity is close to zero). For the saccade is subject to predictions errors and execution noise, the saccade landing position may be different from the initial prediction. The final accuracy, as instantiated in the accuracy map, contains this intrinsic imprecision, and is thus necessary lower than the optimal one. The consequence is that in some cases, the approximate information gain may become negative, when the future accuracy is actually lower than the current one. This is for instance the case when the target is exactly positioned at the center of the fovea.

